# Proteomics study of colorectal cancer and adenomatous polyps identifies TFR1, SAHH, and HV307 as potential biomarkers for screening

**DOI:** 10.1101/2020.10.19.344887

**Authors:** Meifang Tang, Liuhong Zeng, Zhaolei Zeng, Jie Liu, Jie Yuan, Dongjie Wu, Ying Lu, Jin Zi, Mingzhi Ye

## Abstract

Colorectal cancer (CRC) is a malignant tumour with high morbidity and mortality worldwide. Efficient screening strategies for CRC and pre-cancerous lesions will promote early medical intervention and treatment, thus reducing morbidity and mortality. Proteins are generally considered key biomarkers of cancer. Herein, we performed a quantitative, tissue-original proteomics study in a cohort of ninety patients from pre-cancerous to cancerous conditions by liquid chromatography-tandem mass spectrometry. A total of 134,812 peptides, 8,697 proteins, 2,355 (27.08%) union differentially expressed proteins (DEPs), and 409 shared DEPs (compared with adjacent tissues) were identified. The number of DEPs showed a positive correlation with increasing severity of illness. The union and shared DEPs were both enriched in the KEGG pathway of focal adhesion, metabolism of xenobiotics by cytochrome P450, and drug metabolism – cytochrome P450. Among the 2,355 union DEPs, 32 were selected for identification and validation by multiple reaction monitoring from twenty plasma specimens. Of these, three proteins, transferrin receptor protein 1 (TFR1), adenosylhomocysteinase (SAHH), and immunoglobulin heavy variable 3-7 (HV307), were significantly differentially expressed and displayed the same expression pattern in plasma as observed in the tissue data. In conclusion, TFR1, SAHH, and HV307 may be considered as potential biomarkers for screening of CRC.

**Significance:** CRC is a malignant tumour with high morbidity and mortality worldwide. Efficient screening strategies for CRC and pre-cancerous lesions can play an important role in addressing the issue of high morbidity and mortality. Screening of molecular biomarkers provide a non-invasive, cost-effective and effective approach. Proteins are generally considered key molecular biomarkers of cancer. Our study reports a quantitative proteomics analysis of protein biomarkers for colorectal cancer (CRC) and adenomatous polyps and identifies TFR1, SAHH, and HV307 as potential biomarkers for screening. The research makes a significant contribution to the literature because whereas mass spectrometry-based proteomics research has been widely used for clinical research, its application to clinical translation is lacking as parallel specimens ranging from pre-cancerous to cancerous tissues, according to the degree of disease progression, have not been readily assessed.

## 1. Introduction

Colorectal cancer (CRC) is a malignant tumour with high morbidity and mortality worldwide [1]. It occurs in the colon or rectum, which is part of the large intestine of the digestive system, and usually begins as non-cancerous polyps in the lining of the intestine. The accumulation of multiple genetic mutations, epigenetic defects, and other factors can be involved in the development of CRC through the “adenoma-carcinoma” pathway requiring an average of 10-15 years [2–4]. The five-year survival rate of CRC declines sharply when the degree of the disease increases, with stage I reaching more than 90% and stage IV being less than 20% [4, 5]. Studies have pointed out that for every 1% increase in the adenoma detection rate, the incidence of colon cancer can be reduced by 3% and the mortality rate by 4% [6]. Therefore, early screening of CRC and precancerous lesions, such as adenomatous polyps, is particularly essential. Colonoscopy, guaiac faecal occult blood test (gFOBT), faecal immunochemical test for haemoglobin (FIT), and the detection of molecular biomarkers are some of the conventional screening methods [7–11]. As the gold standard, colonoscopy is invasive, operator-dependent, and requires bowel preparation and dietary modification, which might cause complications and poor compliance [12]. Although the faecal occult blood test usually uses guaiac (gFOBT) or specific antibodies (FIT) to detect the presence and concentration of haemoglobin and is easy and cost-effective, the sensitivity for CRC (~79%) varies based on interfering factors [13] and the detection rate of adenomatous polyps is extremely low (~23.8%) [14]. The potential of molecular biomarkers has been gradually explored in recent years, including genetic mutations, methylation, and miRNA markers [11, 14–20], with few protein biomarkers applied. Traditional protein antigens such as carcinoembryonic antigen, which are mainly used for prognoses, perform poorly for screening in the general risk population. Therefore, non-invasive, economic, efficient and early biomarkers are still urgently needed for screening strategies. In recent years, there has been an increasing interest in translational proteomics to understand the mechanism and characteristics of CRC and to provide a good direction for the discovery and validation of protein biomarkers [21]. The types of specimens used include freshly frozen colorectal cancer tissue, formalin-fixed paraffin-embedded tissues, normal or inflamed mucosa, and blood including serum and plasma, among others. The identified protein biomarkers include beta-like 2, dipeptidase 1, olfactomedin-4 (OLFM4), kininogen-1 (KNG1), and transport protein Sec24C (Sec24C). Despite the explosive growth of mass spectrometry-based proteomics research and protein markers, this approach has not been successfully applied to clinical translation, possibly due to the lack of verification of large clinical sample cohorts. One of the greatest challenges is that the protein landscape and expression patterns have rarely been assessed using parallel specimens from pre-cancerous and cancerous conditions according to the degree of disease progression. We thus performed proteomics using specimens ranging from precancerous to cancerous tissues by liquid chromatography-tandem mass spectrometry (LC-MS/MS), which uses data-independent acquisition (DIA) mode for label-free protein quantification and to identify significantly differentially expressed proteins (DEPs) and then verified plasma specimens using liquid chromatography multiple reaction monitoring mass spectrometry (MRM) technology (Fig 1).

**Fig 1.**
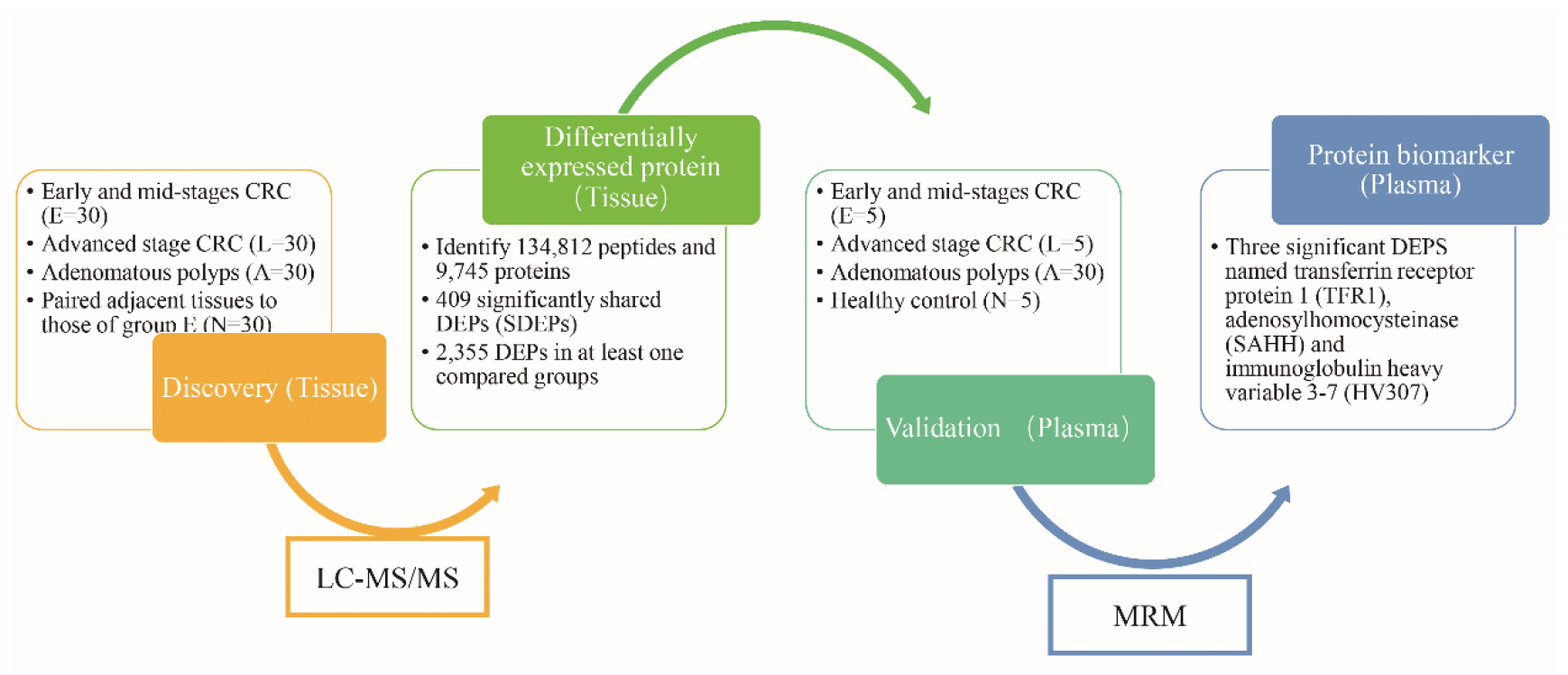
The workflow of the discovery of protein biomarkers with an liquid chromatography-tandem mass spectrometry (LC-MS/MS)-based proteomics landscape to select differentially expressed proteins (DEPs) and MRM-based validation from colorectal cancer (CRC) and adenomatous polyps. Tissue specimens from early and mid-CRC stages (group E, stage I and II), advanced CRC stage (group L, stage III and IV), adenomatous polyps (group A), and the paired paracancerous tissues of group E (group N), with 30 cases in each group, were assessed by LC-MS/MS, which uses data-independent acquisition (DIA) mode for label-free protein quantification and significant DEP selection. A total of 134,812 peptides, 8,697 proteins, 2,355 (27.08%) DEPs, 409 (4.70%) shared DEPs (SDEPs), and 2,355 total DEPs were identified in at least one compared group when compared to the levels in group N. Next, the 32 selected DEPs were verified by multiple reaction monitoring mass spectrometry (MRM) technology using plasma specimens from early and mid-CRC stages (group E, stage I), advanced CRC stage (group L, stage III and IV), adenomatous polyps (group A), and the healthy control tissue (group N) with five cases in each group. A total of three significant DEPs named transferrin receptor protein 1 (TFR1), adenosylhomocysteinase (SAHH), and immunoglobulin heavy variable 3-7 (HV307) were identified as potential protein biomarkers in both tissues and plasma specimens.

## 2. Materials and Methods

### 2.1 Participants

For the DIA analysis, tissue samples from 90 enrolled cases (Table 1) were collected from 2016 to 2017, including 60 cases of CRC and 30 cases of adenomatous polyps in the non-progressive stage. For MRM analysis, plasma samples from another 20 enrolled cases were used, including five CRC cases of stage I, five CRC cases of stage IV, five cases of adenomatous polyps, and five cases of healthy controls (Table S1). All participants were confirmed by pathological biopsy and noticed the experimental protocol.

**Table 1.**
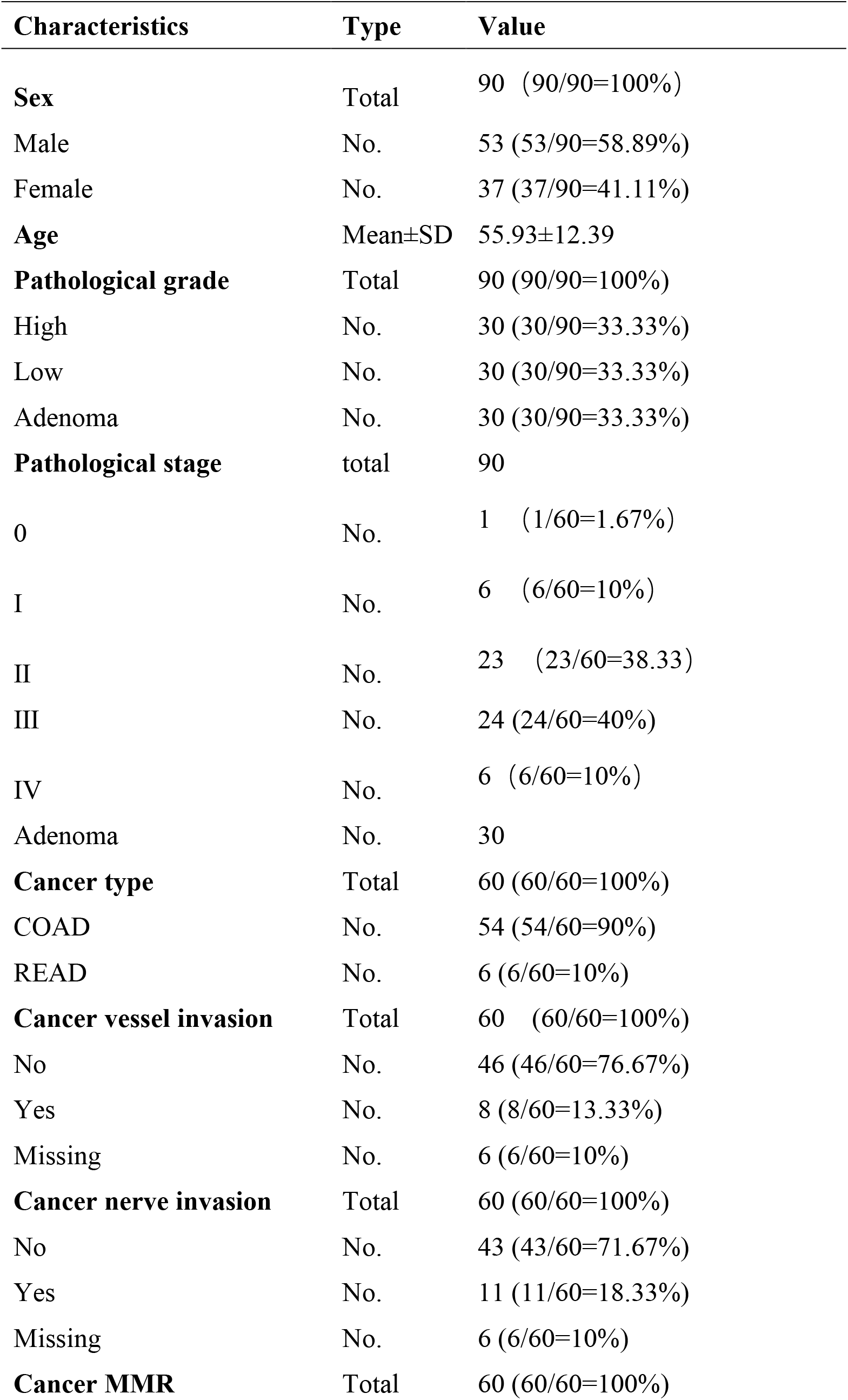
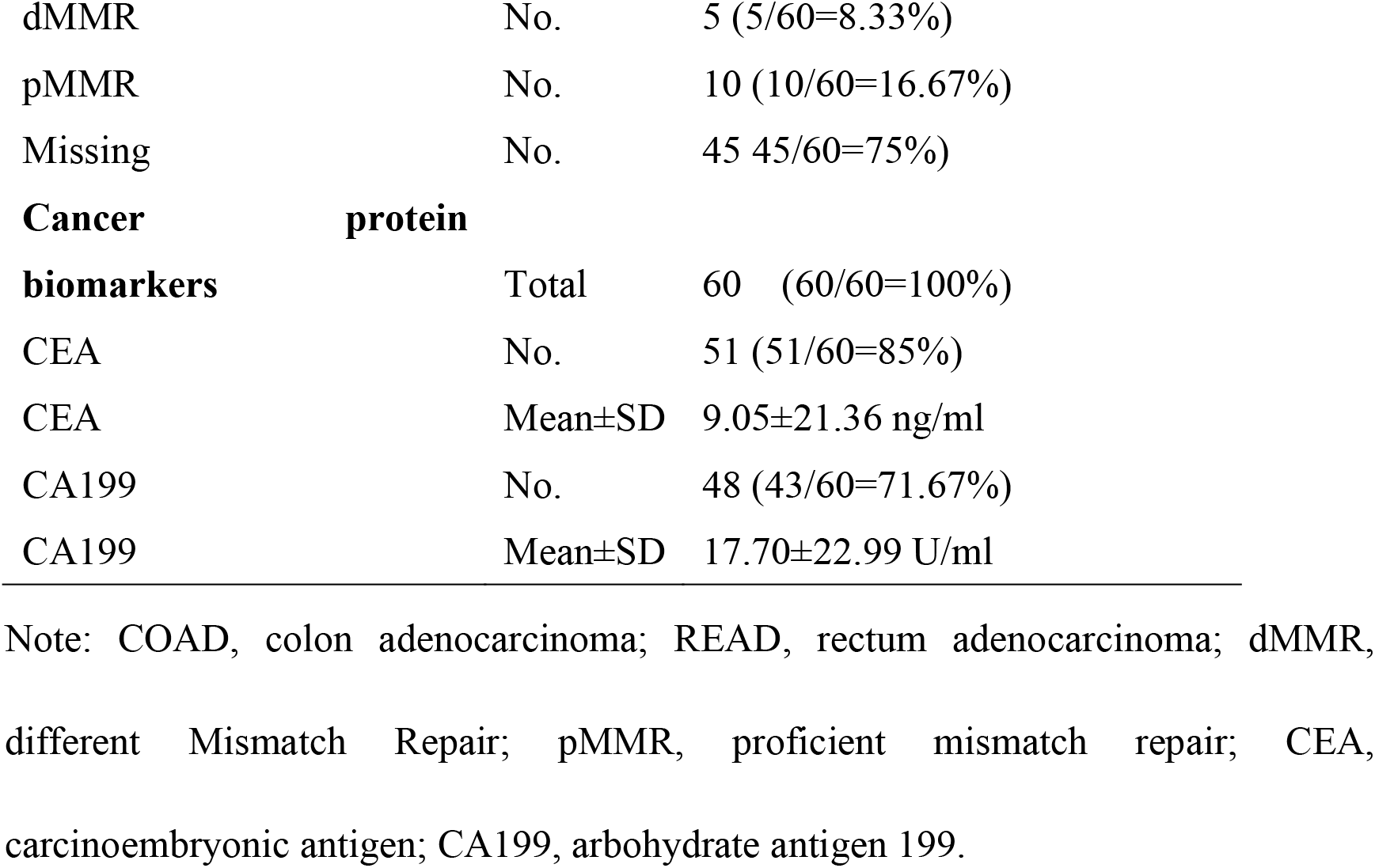
Characteristics and clinical information of 90 enrolled colorectal cancer and adenomatous polyps cases

### 2.2. LC-MS/MS procedure for tissue specimens

#### 2.2.1. Protein Extraction

Samples were ground into powder in liquid nitrogen and then extracted with lysis buffer (7 M urea, 2 M thiourea, 4% CHAPS, 40 mM tris-HCl, pH 8.5) containing 1mM phenylmethylsulfonyl fluoride and 2 mM ethylenediaminetetraacetic acid. Next, 10 mM dithiothreitol (DTT) was added, and after 5 min, the suspension was sonicated at 200 W for 15 min and then centrifuged at 4 °C and 30 000 × *g* for 15 min. The supernatant was mixed well with a 5× volume of chilled acetone containing 10% (v/v) trichloroacetic acid and incubated at −20 °C overnight. After centrifugation at 4 °C and 30 000 × *g*, the supernatant was discarded. Then, the precipitate was washed with chilled acetone three times. The pellet was air-dried and dissolved in lysis buffer (7 M urea, 2 M thiourea, 4% NP40, 20 mM tris-HCl, pH 8.0–8.5). The suspension was sonicated at 200 W for 15 min and centrifuged at 4 °C and 30 000 × *g* for 15 min, and the supernatant was transferred to another tube. To reduce disulphide bonds in proteins of the supernatant, 10 mM DTT was added and incubated at room temperature for 1 h. Subsequently, 55 mM IAM was added to block the cysteines and incubated for 1 h in the dark. Proteins were quantified using a Bradford kit (Bio-Rad, Hercules, CA, USA)..

#### 2.2.2. Protein Digestion

A total of 100 μg protein was digested with Trypsin Gold (Promega, Madison, WI, USA) with the ratio of protein:trypsin = 30:1 at 37 °C for 16 h. After trypsin digestion, the peptides were dried by vacuum centrifugation. Peptides were reconstituted in 0.5 M TEAB.

#### 2.2.3. High pH RP separation

Ten micrograms of peptides from each sample was mixed and 200 μg of the mixed samples was diluted with 2 mL mobile phase A (5% acetonitrile (ACN), pH 9.8), separated using the LC-20AB liquid chromatography (LC) system (SHIMADZU, Kyoto, Japan) with a column (5 μm, 4.6 × 250 mm, Gemini C18) at a flow rate gradient of 1 ml/min. The peptides were eluted with the following linear gradient steps: 5% mobile phase B (95% CAN, pH 9.8) for 10 min, 5% to 35% mobile phase B for 40 min, 35% to 95% mobile phase B for 1 min, mobile Phase B 3 min, and 5% mobile phase B equilibrated for 10 min. The separated peptides were combined to obtain 10 components and then freeze-dried.

#### 2.2.4. High-performance liquid chromatography (HPLC) before mass analysis

The dried peptide samples were reconstituted with mobile phase A (2% ACN, 0.1% FA), centrifuged at 20,000 × *g* for 10 min, and the supernatant was injected into the UltiMate 3000 UHPLC system (Thermo Fisher Scientific, San Jose, CA). The sample was first enriched and desalted by the trap column and then separated using a self-packed C18 column (inner diameter: 150 μm, column material particle size: 1.8 μm, column length: 25 cm), by the following effective gradient at a flow rate of 500 nl/min: 0–5 min, 5% mobile phase B (98% ACN, 0.1% FA); 5–160 min, mobile phase B linearly increased from 5% to 35%; 160≠170 min, mobile phase B increased from 35% to 80%; 170–175min, 80% mobile phase B; 176–180min, 5% mobile phase B. The nanolitre separation was is directly connected to the mass spectrometer.

#### 2.2.5. LC-MS/MS with data-dependent acquisition (DDA) mode for library construction

The peptides separated by HPLC from mixed samples were ionised by the nanoESI source and then analysed using Q-Exactive HF (Thermo Fisher Scientific, San Jose, CA) in DDA mode. The main parameter settings were listed with 1.6 kV of the ion source voltage, 350–1500 m/z of the first-level MS, 100 m/z, 15,000 resolution of the second-level MS, 2+ to 7+ of the charge, greater than 10,000 ranks among the top 20 precursor ions from the screening conditions for the secondary fragmented precursor ions, high collision dissociation (HCD) of the ion fragmentation mode, Orbitrap of the electrostatic trap for the fragment ions, 30 s of the dynamic exclusion time, and Level 1 3E6, Level 2 1E5 of the AGC.

#### 2.2.6. LC-MS/MS in DIA mode

Each digested sample was ionised by the nanoESI source and then entered into Q-Exactive HF (Thermo Fisher Scientific, San Jose, CA) in DIA mode. The main parameter settings were listed with 1.6 kV of the ion source voltage, 350–1500 m/z of the first-level MS, 100 m/z, 12,000 resolution of the second-level MS, 350–1500 Da divided into 40 windows for fragmentation and signal acquisition, HCD of the ion fragmentation mode, Orbitrap of the electrostatic trap for the fragment ions, 30 s of the dynamic exclusion time, and Level 1 3E6, Level 2 1E5 of the AGC.

#### 2.2.7. Data analysis for LC-MS/MS

For the library construction, the fractionated DDA data was identified by MaxQuant [22] with detailed parameter settings, shown in Table S2, and then, the mass spectra library was constructed using Spectronaut [23] with FDR ≤ 1% as a key parameter. For large-scale DIA data, Spectronaut was used during the spectral library construction, and the deconvolution extraction and “mProphet” algorithm was used for data analysis and quality control with FDR ≤ 1%. DEPs between different comparison groups were analysed by MSstats [24] with linear mixed-effects models, and significant DEPs were considered with the conditions of fold change ≥ 2 and P-value < 0.05.

Functional annotation was conducted using gene ontology (GO) (37, 38), clusters of orthologous groups (COG) (39), and Kyoto Encyclopaedia of Genes and Genomes (KEGG) pathway (40) analyses. The following functional enrichment analysis was implemented using the clusterProfiler [25] package with over-representation analysis mode (ORA) [26] and gene set enrichment analysis mode (GSEA) [27]. WoLF PSORT [28] was used to predict the subcellular localisation of the protein. STRING (version:11.0) [29] software was used to analyse functional protein association networks. The active interaction sources came from experiments and databases, the minimum required interaction score was 0.9, and no more than 10 interactors were shown at the 1st shell.

### 2.3. MRM procedure for plasma specimens

Ten microlitres of plasma was diluted using 500 μl lysis buffer as described previously herein. After sonication and centrifugation, protein quantitation in the supernatant was determined using the method described. For protein digestion, 50 fmol of β-galactosidase was added to each 100 μg sample and then subjected to the same digestion protocol as described previously herein.

#### 2.3.1 MRM

A QTRAP 6500 mass spectrometer (SCIEX, Framingham, MA, USA) equipped with an LC-20AD nano HPLC system (Shimadzu, Kyoto, Japan) was used for MRM analysis. The mobile phase consisted of solvent A, 0.1% aqueous formic acid, and solvent B, 98% acetonitrile with 0.1% formic acid. Peptides were separated on a C18 column (0.075 × 150 mm column, 3.6 μm) at 300 nl/min and eluted with a gradient of 5-30% solvent B for 38 min and 30–80% solvent B for 4 min and maintained at 80% for 8 min. For the QTRAP 6500 mass spectrometer, a spray voltage of 2400 V, nebuliser gas of 23 p.s.i., and a dwell time of 10 ms were used. Transitions for each peptide were monitored using unit resolution in both Q1 and Q3 quadrupoles to maximise specificity (Table S3).

#### 2.3.2 Data analysis for MRM

Skyline [30] was used to integrate the raw file generated by QTRAP 6500. An iRT strategy was used to define the chromatography of a given peptide against a spectral library. All transitions for each peptide were used for quantitation unless interference from the matrix was observed. A spiked β-galactosidase was used for label-free data normalisation. MSstats [24] with the linear mixed-effects model were calculated for DEPs, and the P-values were adjusted to control the FDR at a cut-off of 0.05. All DEPs with an FDR < 0.05 and fold-change no less than 1.4 were considered significant.

### 2.4. Statistical analysis

All statistical analysis was performed and graphs were generated using R (version 3.50). The box plot was produced using the ggboxplot function; analysis of variance was used to analyse the differences among the four groups. The receiver operating characteristic curve [31] analysis was implemented using the pROC [32] package.

## 3. Result

### 3.1. Proteomics landscape of CRC and adenomatous polyps

In DDA mode, a total of 121,937 peptides and 9,745 proteins were identified for library construction. Among the identified proteins (details for the identified proteins are listed in Table S4 and shown in Fig. S1-A), 9,008 proteins contained at least two unique peptides (mean, 11.59; range, 1–249; 95% CI, 11.31–11.86; 95th-percentile, 34), accounting for 92.44% of the total. A total of 5,778 proteins with a sequence coverage of at least 20% (mean, 28.34%, range, 2–100%; 95% CI, 27.96–28.73%; 95th-percentile, 65.17%) accounted for 59.29%.

In DIA mode, a total of 134,812 peptides (FDR < 1%, Table S4) and 8,697 proteins (Table S5, Table S6, Table S7) were identified. The coefficient of variation (CV) of quantitative values both from peptides and proteins in each group was less than 10% (Fig. S2-A), revealing that the experiments of label-free protein quantification remained steady and repeatable. With an increasing severity of illness, a slight increase in the number of peptides (Fig. S3) and proteins (Fig. 1-A) was recorded. Principal component analysis of protein quantitative values from each sample in each group (Fig. 2-B, Fig. S2-B) showed a clear distinction between group N and the remaining three groups, especially for group L. However, there was no obvious distinction between adenomatous polyps (group A) and CRC (group E or L). A possible explanation for this might be that altered expression based on the number of proteins appeared even at the pre-cancer stage. Meanwhile, the heatmap diagram of scale values from quantitation with all identified proteins (Fig. 2-C, Table S7) indicated that the expression patterns between groups E and L were similar. Thrre was significant differences between groups N and CRC (group E or L) and between groups A and group N or CRC (group E or L). Fuzzy clustering (Fig. S4, Table S8) of time series analysis with mean quantitative values in each group from the total proteins was performed with 26 clusters (error = 0.013). Compared to group N, the clusters with an overexpression tendency are listed as clusters 3, 11, 14, and 18, and with those having an underexpression tendency listed as cluster 13. The overexpression tendency clusters were enriched in KEGG pathways of RNA transport, spliceosome, ribosome biogenesis in eukaryotes, aminoacyl-tRNA biosynthesis, proteasome, mRNA surveillance pathway, nucleotide excision repair, RNA degradation, protein processing in endoplasmic reticulum and mismatch repair by ORA analysis, and concerned with the transport, biogenesis, and repair in abnormal cells. Meanwhile, the underexpression tendency clusters were enriched in the KEGG pathways of oxidative phosphorylation, thermogenesis, histidine metabolism, drug metabolism – cytochrome P450, tryptophan metabolism, fatty acid degradation, arachidonic acid metabolism, retrograde endocannabinoid signalling, tyrosine metabolism, metabolism of xenobiotics by cytochrome P450, and related to metabolism.

**Fig. 2.**
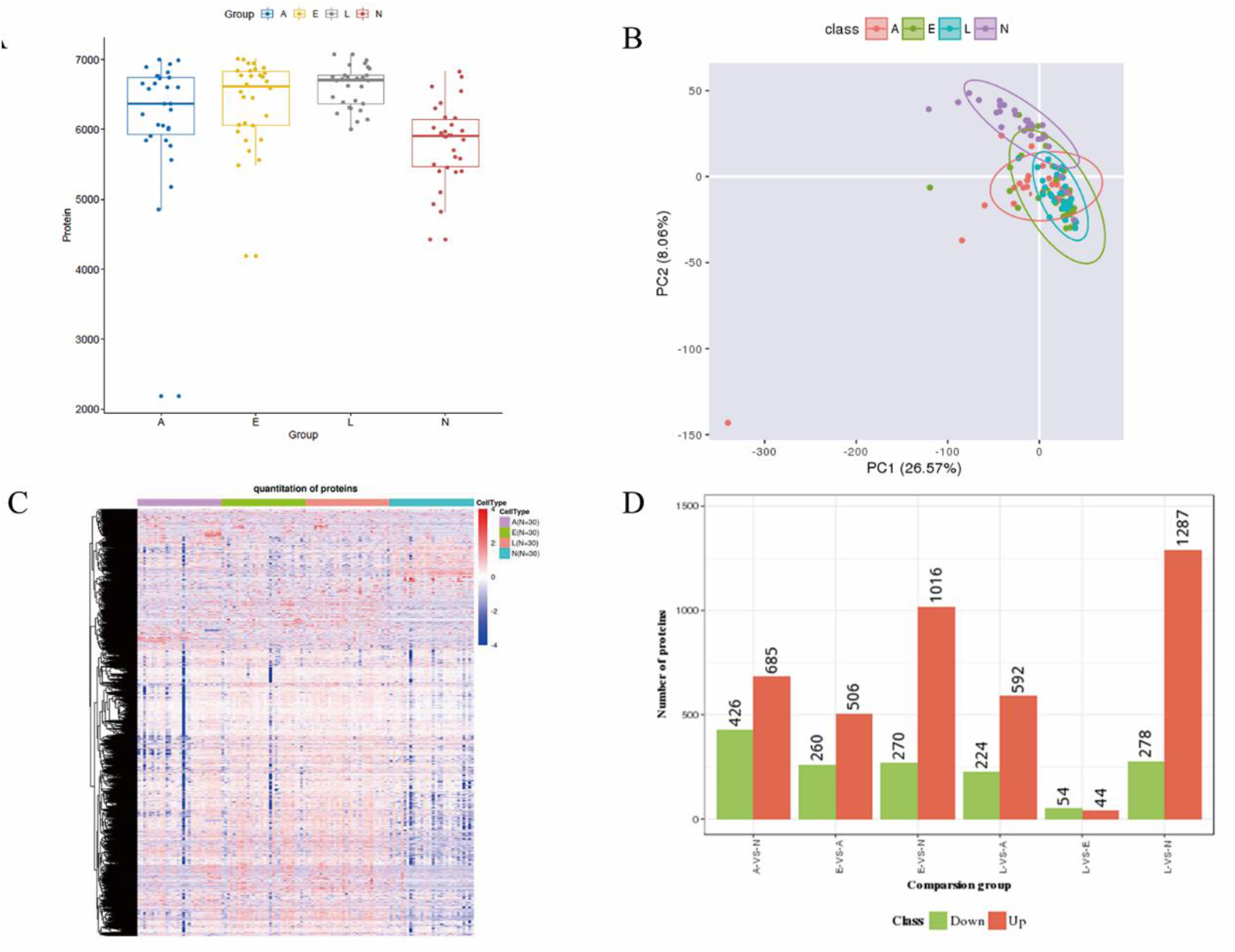
The proteomics landscape of colorectal cancer (CRC) and adenomatous polyps. A. Box plots of the number of identified proteins in group A, E, L, and N. B. Scatter plot of principal component analysis of protein quantitative values from each sample in each group. C. the heatmap diagram of scale values from quantitation with all identified proteins. Each row represents a protein, and each column represents a sample. D. Histogram of the number of significantly differentially expressed proteins (DEPs) in pairwise comparisons (a total of six comparison combinations). The orange colour displays overexpressed DEPs, and the green colour displays underexpressed DEPs. “-VS-” means comparison, the group before VS means the treatment group, and the group after VS means the control group. e.g. “E-VS-N” represents a comparison between group N as the control group and E as the treatment group. Group E, early and midstage CRC: L, advanced stage CRC; A, adenomatous polyps; N, paired adjacent tissues to those of group E.

Compared to those for group A, the clusters with overexpression tendency are listed as clusters 6 and cluster 22, with those having an underexpression tendency are listed as cluster 7. The underexpression tendency clusters were enriched in the KEGG pathways of complement and coagulation cascades, Th1 and Th2 cell differentiation, and phagosomes.

A total of six comparison combinations and 2,752 significant DEPs were produced when compared to any two of the four groups (Fig. 2-D, TableS9). It was apparent from this graph that significant DEPs were predominantly overexpressed except for those discovered by the comparison “L-VS-E”. When compared to those in group N, the numbers of significant DEPs of the comparisons “A-VS-N”, “E-VS-N”, and “L-VS-N” were 1,111 (12.77%), 1,286 (14.79%), and 1,565 (17.99%), respectively. There were 426 (4.9%), 270 (3.10%), and 278 (3.20%) significantly underexpressed DEPs and 685 (7.88%), 1016 (11.68%), and 1287 (14.80%) significantly overexpressed DEPs, respectively. Consistent with these findings, we found that the number of significant DEPs also increased as the severity of disease increased. This result might be explained by the fact that significantly overexpressed DEPs correspondingly increased following the increasing severity of illness. Simultaneously, when compared to that in group A, the numbers of significant DEPs of the comparisons “N-VS-A”, “E-VS-A”, and “L-VS-A” were 1,111 (12.77%), 766 (8.81%), and 816 (9.38%), respectively. There were 685 (7.88%), 260 (2.99%), and 224 (2.58%) significantly underexpressed DEPs and 426 (4.9%), 506 (5.82%), and 592 (6.81%) significantly overexpressed DEPs, respectively. Interestingly, a greater difference in “N-VS-A” was observed compared to that with in “E-VS-A” or “L-VS-A”. In addition, the number of DEPs fell to a low value of 98 (1.13%) for “L-VS-E”.

### 3.2. Pathway analysis of significant DEPs between CRC and adenomatous polyps

The next section of the research was concerned based on the significant DEPs and their functional enrichment analysis when compared to group N. There were 409 (4.70%; Fig. 3-A) significant shared DEPs (SDEPs) for comparisons “A-VS-N”, “E-VS-N”, and “L-VS-N” with 282 overexpressed SDEPs (3.24%) and 127 underexpressed SDEPs (1.46%). Further, 68.95% of those were overexpressed, implying that protein overexpression might play an important role in the development of adenomatous polyps and CRC. The heatmap diagram of scale values from quantitation with 409 SDEPS indicated that the expression pattern between adenomatous polyps (group A) and CRC (group E or L) was strikingly similar. However, dramatic differences in the expression patterns between group N and the other groups were found. Next, The SDEPs were subjected to GO (Fig. S5) and KEGG pathway (Fig. 3-C) functional enrichment analysis by GSEA based on the log2 fold-change value from the “E-VS-N” comparison. SDEPs were enriched in GO terms organelle part (including intracellular membrane-bounded organelle and intracellular organelle part), nucleus and gene expression, and ECM-receptor interaction, focal adhesion, drug metabolism – cytochrome P450, and drug metabolism – cytochrome P450 (Table S10) pathways. Concerns were expressed regarding disturbances in signal transduction, transport, and metabolism with respect to SDEPS in the cells between CRC and adenomatous polyps. All proteins related to these pathways, including integrin alpha-9, collagen alpha-2(VI) chain, laminin subunit alpha-5, collagen alpha-1(I) chain, platelet glycoprotein 4, laminin subunit beta-2, tenascin-X, caveolin-1, myosin regulatory light polypeptide 9, filamin-C, glutathione S-transferase A1, UDP-glucuronosyltransferase 2A1, UDP-glucuronosyltransferase 2A3, glutathione S-transferase Mu 2, alcohol dehydrogenase 1C, and alcohol dehydrogenase 1B, were significantly underexpressed in the CRC and adenomatous polyps groups. In addition, the results of the protein interaction network of 409 SDEPs (confidence ≥ 0.9) showed that the top participating nodes were NHP2L1, SKIV2L2, CCNA2, U2AF1L4, and SRSF6 (Fig. 3-D, Table S11).

**Fig. 3.**
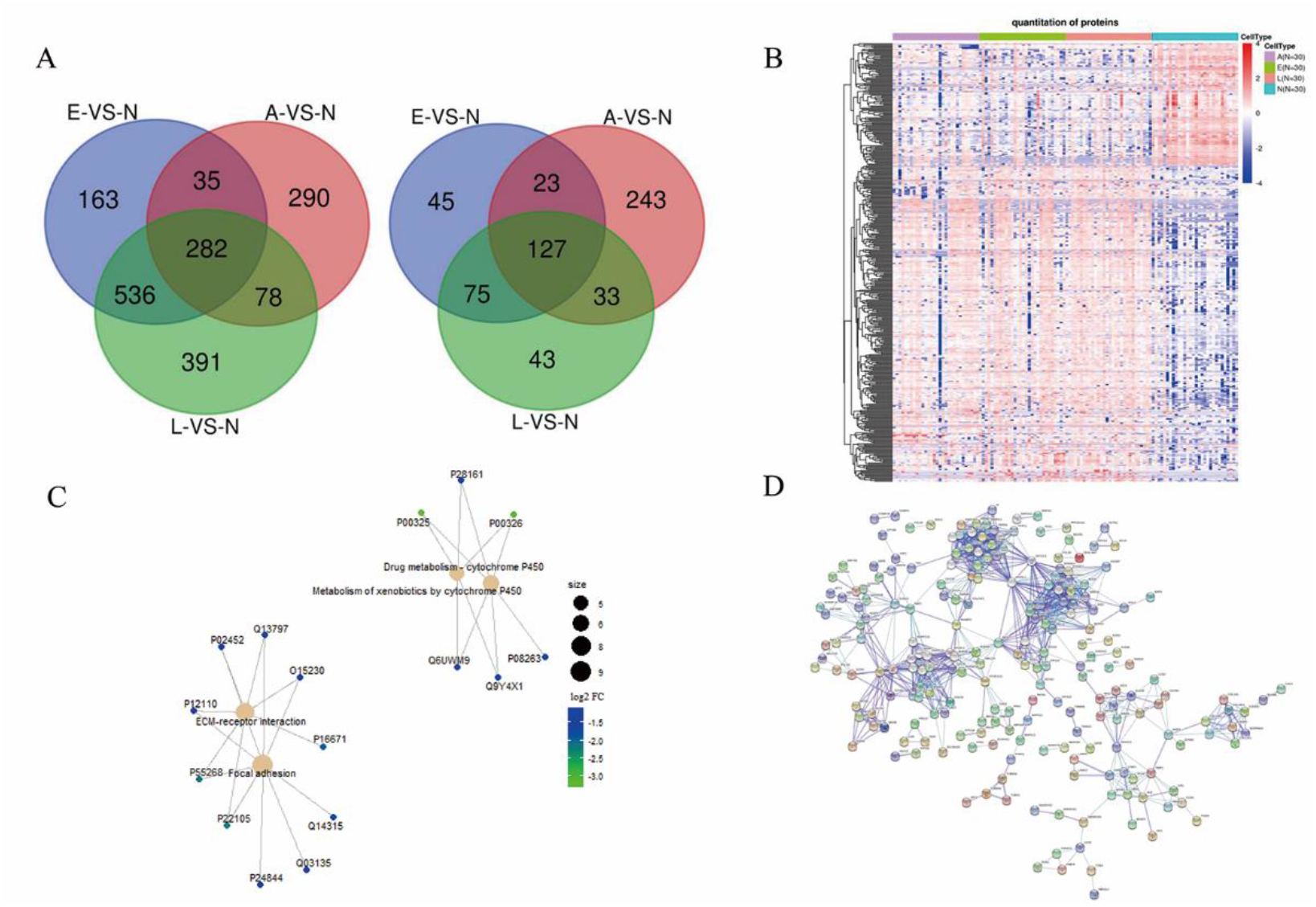
Significant shared differentially expressed protein (SDEP) landscape of colorectal cancer (CRC) and adenomatous polyps. A. Venn diagram of the number of DEPs among the three comparison groups, namely “A-VS-N”, “E-VS-N”, and “L-VS-N”) with 282 significantly overexpressed DEPs (left) and 127 significantly underexpressed DEPs (right). B. Heatmap diagram of 409 SDEPs in each sample from each group showing that there was a clear distinction between group N and the other groups. Each row represents a protein, and each column represents a sample. C. KEGG pathway functional enrichment network of 409 SDEPs by gene set enrichment analysis mode (GSEA) with an input value of log2 fold-change in the “E-VS-N” comparison. D. The protein interaction network of 409 SDEPs (confidence ≥ 0.9).

However, there were 2,355 (27.08%) significant union DEPs in total that were differentially expressed in at least one group when compared to levels in group N, specifically “A-VS-N”, “E-VS-N”, and “L-VS-N”. Functional enrichment analysis with GSEA (FDR < 0.05, Fig. 4-A) for the 2,355 proteins with the input value being log2 fold-change for the comparison “E-VS-N” showed that DEPs were enriched in KEGG pathways of DNA replication, renin secretion, tyrosine metabolism, steroid hormone biosynthesis, bile secretion, alcoholism, retinol metabolism, neuroactive ligand-receptor interaction, mineral absorption, cell adhesion molecules, metabolism of xenobiotics by cytochrome P450, drug metabolism – cytochrome P450, pancreatic secretion, complement and coagulation cascades, and focal adhesion (Table S11). All proteins related to the DNA replication pathways, including replication licencing factors such as MCM2, MCM3, MCM4, MCM5, MCM6, and MCM7, replication factor C subunits such as RFC1, RFC2, RFC3, RFC4, and RFC5, DNA polymerase alpha catalytic subunit, DNA polymerase epsilon subunit 3, DNA primase large subunit, DNA polymerase delta subunit 3, and proliferating cell nuclear antigen were significantly overexpressed in groups E and L. In addition, other enriched pathways were significantly underexpressed in groups E and L. These findings suggested that the roles of these pathways were associated with the control of proliferation, transport, and metabolism of tumour cells.

### 3.3. Validation of SDEPs between CRC and adenomatous polyps

Plasma specimens were used for verification because the collection of plasma samples is non-invasive, acquisition is user-friendly, and this is suitable for colorectal cancer diagnosis and screening for high-risk populations. All 2,355 DEPs were subjected to MRM method development and only 32 target proteins including 11 SDEPs could be identified by MRM from plasma specimens (Table S3). The results from the MRM platform (Table S13-S15) showed that there were 13 significant DEPs (accounting for 40.625%) in plasma samples, and three of them showed the same expression patterns between tissue and plasma samples in the four groups (Fig. 5). The proteins were transferrin receptor protein 1 (TFR1, P02786), adenosylhomocysteinase (SAHH, P23526), and immunoglobulin heavy variable 3-7 (HV307, P01780). The first two proteins were significantly overexpressed proteins and the last was significantly underexpressed in groups A, E, and L when compared to levels in group N. Of note, based on the figure, strong consistency in the expression patterns from the three DEPs between tissue specimens and plasma specimens was observed. Next, the performance of the area under curve (AUC) in each comparison group of these three proteins from tissue and plasma specimens was also assessed, as shown in Table 2. The AUCs of TFR1 (P02786) from plasma samples in “L-VS-N”, “E-VS-N”, “A-VS-N” and “E-VS-A” comparisons were 0.960, 0.960, 0.800, and 0.840, respectively. Simultaneously, the AUCs of SAHH (P23526) from tissue samples in “L-VS-N”, “E-VS-N”, and “E-VS-A” comparisons were 0.938, 0.896, and 0.920, respectively, and those from plasma samples of “L-VS-N”, “E-VS-N”, and “A-VS-N” comparisons were 0.920, 1, and 0.840. Further, the AUCs of HV307 (P01780) from tissue samples in the “L-VS-N”, “E-VS-N”, and “E-VS-A” comparisons were 0.925, 0.887, and 0.834, respectively, and those from plasma samples of “L-VS-N” and “E-VS-N” comparisons reached 1 and 0.880, respectively. The AUC values listed in red font were all no less than 0.8, demonstrating that TFR1 (P02786), SAHH (P23526), and HV307 (P01780) might be potential biomarkers, in both tissue and plasma samples, for CRC and adenomatous polyps.

**Fig. 4.**
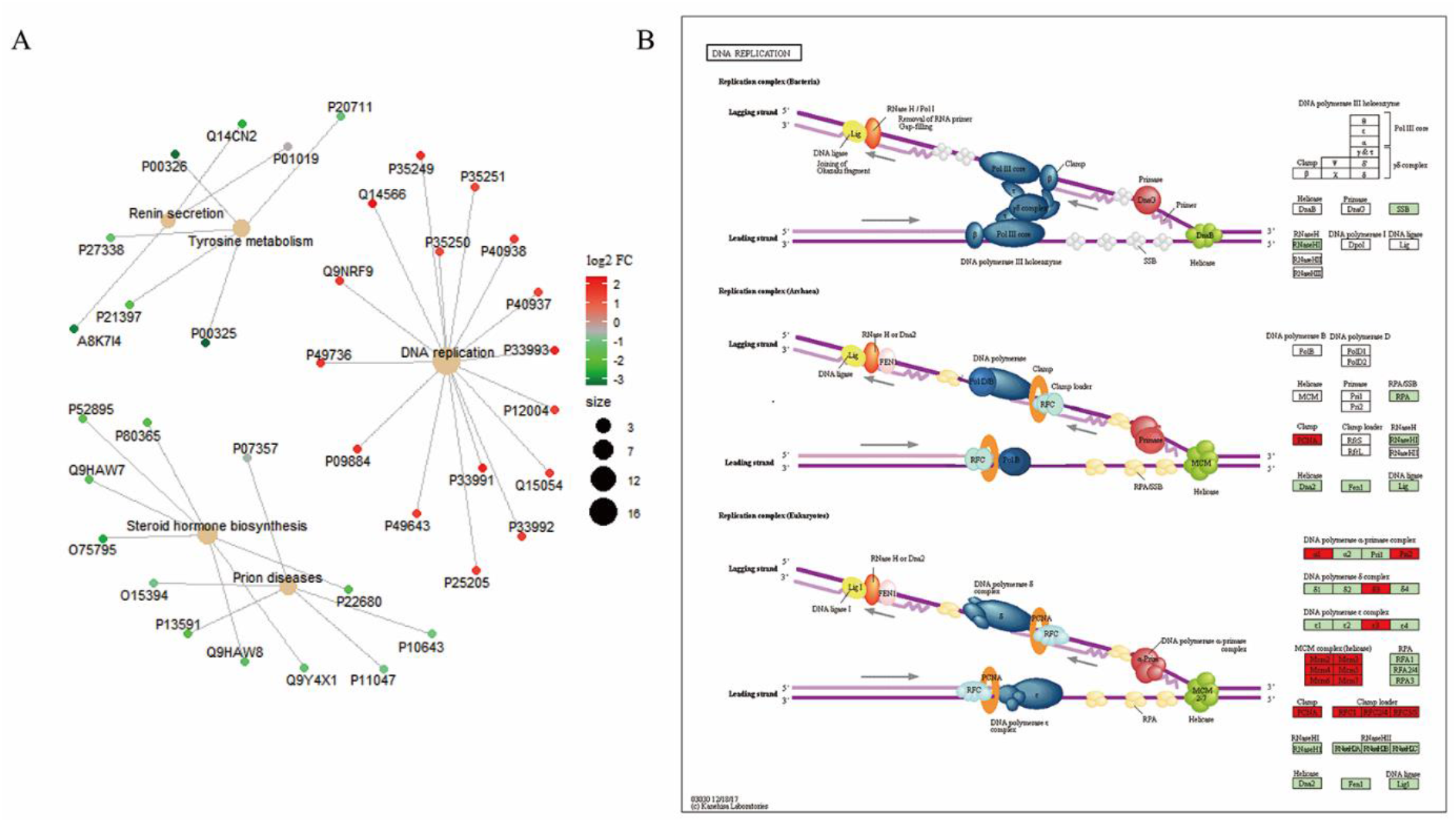
Pathway analysis of the 2,355 significant differentially expressed proteins (DEPs). A. KEGG pathway functional enrichment network of 2,355 DEPs by gene set enrichment analysis mode (GSEA) in the “E-VS-N” comparison. B. KEGG pathway of DNA replication. Sixteen proteins related to the DNA replication pathways were completely significantly overexpressed in colorectal cancer (CRC) patients and partly significantly overexpressed in adenomatous polyps patients.

**Fig. 5.**
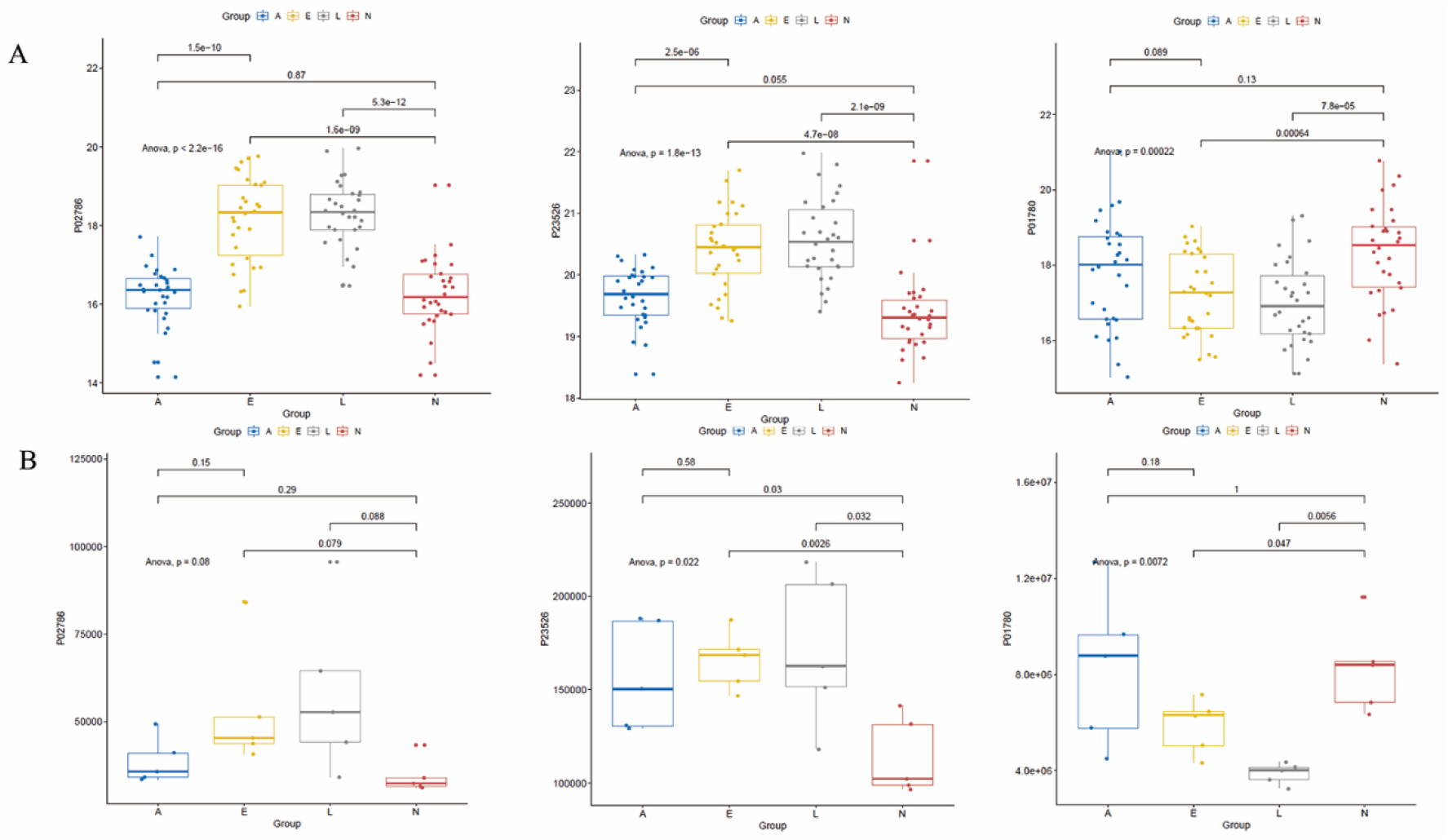
Boxplot of the abundance of TFR1 (P02786), SAHH (P23526), and HV307 (P01780) in the colorectal cancer (CRC), adenoma, and control groups from tissue (A) and plasma (B) specimens. The vertical coordinate of A is the log2 expression, and the vertical coordinate of B is the expression.

**Table 2.** Area under the curve (AUC) of the performance of different comparison groups of tissue samples by LC-MS/MS and of plasma samples by multiple reaction monitoring (MRM).

| Tissue by LC-MS/MS | AUC (95%CI) |  |  |
| --- | --- | --- | --- |
| Comparison | TFR1 (P02786) | SAHH (P23526) | HV307 (P01780) |
| L-VS-N | 0.786 (95%CI, 0.668–0.903) | 0.938 (95%CI, 0.872–1.000) | 0.925 (95%CI, 0.85–1.000) |
| E-VS-N | 0.751 (95%CI, 0.627–0.875) | 0.896 (95%CI, 0.816–0.975) | 0.887 (95%CI, 0.797–0.977) |
| A-VS-N | 0.617 (95%CI, 0.473–0.761) | 0.509 (95%CI, 0.358–0.660) | 0.710 (95%CI, 0.575–0.845) |
| E-VS-A | 0.623 (95%CI, 0.478–0.768) | 0.920 (95%CI, 0.847–0.993) | 0.834 (95%CI, 0.728–0.941) |
| Plasma by MRM | Protein AUC (95%CI) |  |  |
| Comparison | TFR1 (P02786) | SAHH (P23526) | HV307 (P01780) |
| L-VS-N | 0.960 (95%CI, 0.849–1.000) | 0.920 (95%CI, 0.736–1.000) | 1.000 (95%CI, 1.000–1.000) |
| E-VS-N | 0.960 (95%CI, 0.849–1.000) | 1.000 (95%CI, 1.000–1.000) | 0.880 (95%CI, 0.658–1.000) |
| A-VS-N | 0.800 (95%CI, 0.472–1.000) | 0.840 (95%CI, 0.568–1.000) | 0.520 (95%CI, 0.070–0.970) |
| E-VS-A | 0.840 (95%CI, 0.568–1.000) | 0.600 (95%CI, 0.171–1.000) | 0.720 (95%CI, 0.344–1.000) |

## 4. Discussion

Protein biomarkers for screening have been reported for tissue and blood samples based on the disease type of CRC and adenomatous polyps. However, most research on protein markers is limited in that it has focused on one stage, specifically cancer progression or the precancerous period, such as adenomatous polyps, and has seldomly utilised specimens from pre-cancerous to cancerous conditions according to the degree of disease progression in parallel. At the same time, most protein biomarkers screened in previous research have not been applied clinically. Proteomics studies of CRC and the pre-cancerous period, such as adenomatous polyps, are thus urgently needed. Our study pioneered the use of tissue-original proteomics profiles of CRC with various stages and non-progressive adenomatous polyps, which provides a deeper insight into expression patterns at the protein level from pre-cancerous to cancerous stages, discovering protein biomarkers for diagnosis or screening.

We identified a total of 134,812 peptides and 8,697 proteins, which is slightly more than a previous study (FDR < 1%) [33]. The CV of quantitative values from both peptides and proteins and the scatter plot of quality control samples clustered in a concentrated manner in a principal component analysis (Fig. S2-A), revealing the stability, reliability, and repeatability of the experiments with label-free protein quantification. It is interesting to note that there was a slight increase in the numbers of peptides, proteins, and DEPs with an increasing severity of illness. This result might be explained by the fact that disturbances in protein expression are exacerbated as the degree of disease increases. The insights gained from fuzzy clustering of time series analysis at the protein level could also be of assistance in understanding the development of CRC. The principal component and heatmap analyses of protein quantitative values have extended our knowledge, indicating that there is a clear distinction in the protein expression pattern between paracancerous tissue and CRC or adenomatous polyp tissue. The results of this study support the idea that it is possible to discover protein biomarkers for the diagnosis and screening of CRC and adenomatous polyps. Some protein expression patterns in adenomatous polyps were similar to those of colorectal cancer, whereas some were similar to those of paracancerous tissues. One interesting finding is that in general, the protein expression pattern of adenomatous polyps seems to be more similar to that of colorectal cancer. These relationships might be partly explained by altered expression at the protein level during the pre-cancerous period.

There were 409 (4.70%) SDEPs, and a total of 2,355 union DEPs were identified in the “A-VS-N”, “E-VS-N”, and “L-VS-N” comparisons. Closer inspection of the DEPs table revealed that some traditional CRC oncogenes such as KRAS and NRAS or suppressor genes such as APC and PTEN, which might be frequently mutated at the genomic level, were not significantly altered at the protein expression level. The implication of this is the possibility of personalised diagnosis and screening at the protein level. These SDEPS shared a very similar expression pattern between adenomatous polyps and CRC. In addition, SDEPs and union DEPs shared the same pathways of focal adhesion, metabolism of xenobiotics by cytochrome P450, and drug metabolism – cytochrome P450. The KEGG pathways in which 2,355 enriched proteins were underexpressed in CRC patients, except for DNA replication, were associated with the control of proliferation, transport, and metabolism of tumour cells. Interestingly, focal adhesion, one of the KEGG pathways associated with the enrichment of SDEPs or union DEPs, was significantly underexpressed in the CRC and adenomatous polyps groups; this has been relatively popular in the field of cancer-related signalling mechanisms and pathways in recent years and has been reported to be altered in various type of cancers including ovarian cancer, lung cancer, breast cancer, and gastric cancer [34–37].

From tissue discovery by LC-MS/MS to plasma verification by MRM, there were not many significant DEPs, which could be mainly due to the differences in sample types from tissue to plasma, as the concentration of protein biomarkers in the blood would be too low to detect in early stages and with precancerous lesions. A total of three significant DEPs, namely TFR1, SAHH, and HV307 were identified as potential protein biomarkers in both tissues and plasma specimens. TFR1, encoded by the *TFRC* gene, belongs to the peptidase M28 family and M28B subfamily. The signal-anchor, transmembrane, and transmembrane helix domain is required for iron import from transferrin into cells via endocytosis. Recent studies have reported that TFR1 is involved in kinds of diseases, including cancers, anaemia, and neurodegenerative diseases. Most importantly, TFR1 was verified to be abnormally expressed in various cancers, and some experimental and clinical drugs and antibodies targeting this protein have shown strong anti-tumour effects [38–46]; herein, TFR1 is suggested to be a potential molecular biomarker for targeted cancer diagnosis and therapy. SAHH, encoded by the *AHCY* gene, belongs to the adenosylhomocysteinase family and is a competitive inhibitor of S-adenosyl-L-methionine-dependent methyltransferase reactions and might play a key role in the control of methylation by regulating the intracellular concentration of adenosylhomocysteine. It has been previously reported to be involved in oesophageal squamous cell carcinoma [47] and CRC [48]. Finally, HV307, encoded by the *IGHV3-7* gene, is a V region of the variable domain of immunoglobulin heavy chains that participate in antigen recognition and has been reported to be associated with immune system diseases, lymphoma, and leukaemia. The AUC values of these three proteins for some comparisons reached at least 0.8, and some even reached 1, and thus, TFR1, SAHH, and HV307 might be crucial protein biomarkers for CRC. Moreover, TFR1 and SAHH might be competitive protein biomarkers for adenomatous polyps. The performance of these three potential proteins needs to be verified by clinical validation experiments based on larger blood sample cohorts and making comparisons to the current screening methods.

## 5. Conclusion

The investigation of quantitative proteomics analysis using parallel specimens ranging from pre-cancerous to cancerous tissues, according to the degree of disease progression, has significant implication for the understanding of the development of CRC at the protein level. Our result also has shown that TFR1, SAHH, and HV307 may be considered as candidate protein biomarkers for screening of CRC and pre-cancerous lesions.

## Author contributions

Meifang Tang: Conceptualization, Methodology, Supervision, Writing – Review & Editing. Liuhong Zeng: Data curation, Formal analysis, Visualization, Writing-Original draft preparation. Zhaolei Zeng: Conceptualization, Methodology, Resources. Jie Liu: Software, Validation. Jie Yuan: Resources, Investigation. Dongjie Wu: Investigation. Ying Lu: Investigation. Jin Zi: Writing – Review & Editing. Mingzhi Ye: Funding acquisition, Resources.

## Compliance with ethical standards

The study was approved by appropriate Institutional Review Boards (IRB) of the BGI (NO. BGI-IRB15100-T1).

## Acknowledgements

This research was supported by the Guangzhou Science and Technology Plan Projects (Health Medical Collaborative Innovation Program of Guangzhou) (grant No. 201803040019, 201400000004-5), Guangzhou Key Laboratory of Cancer Trans-Omics Research (GZ2012, NO348).

## Data resource

The data reported in this study are also available in the CNGB Nucleotide Sequence Archive (CNSA: https://db.cngb.org/cnsa; accession number CNP0001326).

## References

1. Bray F, Ferlay J, Soerjomataram I, Siegel RL, Torre LA, Jemal A. Global cancer statistics 2018: GLOBOCAN estimates of incidence and mortality worldwide for 36 cancers in 185 countries. CA Cancer J Clin 2018;68:394–424.

2. Fearon ER. Molecular genetics of colorectal cancer. Annu Rev Pathol 2011;6:479–507.

3. Vogelstein B, Papadopoulos N, Velculescu VE, Zhou S, Diaz LA, Jr., Kinzler KW. Cancer genome landscapes. Science 2013;339:1546–58.

4. Arvelo F, Sojo F, Cotte C. Biology of colorectal cancer. Ecancermedicalscience. 2015;9:520.

5. Feo L, Polcino M, Nash GM. Resection of the primary tumor in stage IV colorectal cancer: When is it necessary? Surg Clin North Am 2017;97:657–69.

6. Corley DA, Jensen CD, Marks AR, Zhao WK, Lee JK, Doubeni CA, Zauber AG, de Boer J, Fireman BH, Schottinger JE, Quinn VP, Ghai NR, Levin TR, Quesenberry CP. Adenoma detection rate and risk of colorectal cancer and death. N Engl J Med. 2014 Apr 3;370(14):1298–306.

7. Schreuders EH, Ruco A, Rabeneck L, Schoen RE, Sung JJ, Young GP, Kuipers EJ. Colorectal cancer screening: a global overview of existing programmes. Gut 2015;64:1637–49.

8. Issa IA, Noureddine M. Colorectal cancer screening: An updated review of the available options. World J Gastroenterol 2017;23:5086–96.

9. Navarro M, Nicolas A, Ferrandez A, Lanas A. Colorectal cancer population screening programs worldwide in 2016: An update. World J Gastroenterol. 2017;23:3632–42.

10. Provenzale D, Gupta S, Ahnen DJ, Markowitz AJ, Chung DC, Mayer RJ, Regenbogen SE, Blanco AM, Bray T, Cooper G, Early DS, Ford JM, Giardiello FM, Grady W, Hall MJ, Halverson AL, Hamiltion SR, Hampel H, Klapman JB, Larson DW, Lazenby AJ, Llor X, Lynch PM, Mikkelson J, Ness RM, Slavin TP, Sugandha S, Weiss JM, Dwyer MA, Ogba N. NCCN Guidelines Insights: Colorectal Cancer Screening, Version 1.2018. J Natl Compr Canc Netw. 2018;16:939–49.

11. Wolf AMD, Fontham ETH, Church TR, Flowers CR, Guerra CE, LaMonte SJ, Etzioni R, McKenna MT, Oeffinger KC, Shih YT, Walter LC, Andrews KS, Brawley OW, Brooks D, Fedewa SA, Manassaram-Baptiste D, Siegel RL, Wender RC, Smith RA. Colorectal cancer screening for average-risk adults: 2018 guideline update from the American Cancer Society. CA Cancer J Clin 2018;68:250–81.

12. Church J. Complications of colonoscopy. Gastroenterol Clin North Am 2013;42:639–57.

13. Lee JK, Liles EG, Bent S, Levin TR, Corley DA. Accuracy of fecal immunochemical tests for colorectal cancer: systematic review and meta-analysis. Ann Intern Med 2014;160:171.

14. Imperiale TF, Ransohoff DF, Itzkowitz SH, Levin TR, Lavin P, Lidgard GP, Ahlquist, DA, Berger, BM. Multitarget stool DNA testing for colorectal-cancer screening. N Engl J Med 2014;370:1287–97.

15. Ganepola GA, Nizin J, Rutledge JR, Chang DH. Use of blood-based biomarkers for early diagnosis and surveillance of colorectal cancer. World J Gastrointest Oncol 2014;6:83–97.

16. Dickinson BT, Kisiel J, Ahlquist DA, Grady WM. Molecular markers for colorectal cancer screening. Gut 2015;64:1485–94.

17. Peluso G, Incollingo P, Calogero A, Tammaro V, Rupealta N, Chiacchio G, Sandoval Sotelo ML, Minieri G, Pisani A, Riccio E, Sabbatini M, Bracale UM, Dodaro CA, Carlomagno N. Current tissue molecular markers in colorectal cancer: A literature review. BioMed Res Int 2017;2017:1–8.

18. Nikolaou S, Qiu S, Fiorentino F, Rasheed S, Tekkis P, Kontovounisios C. Systematic review of blood diagnostic markers in colorectal cancer. Tech Coloproctol 2018;22:481–98.

19. Toma SC, Ungureanu BS, Patrascu S, Surlin V, Georgescu I. Colorectal cancer biomarkers - A new trend in early diagnosis. Curr Health Sci J 2018;44:140–6.

20. Marcuello M, Vymetalkova V, Neves RPL, Duran-Sanchon S, Vedeld HM, Tham E, van Dalum G, Flügen G, Garcia-Barberan V, Fijneman RJ, Castells A, Vodicka P, Lind GE, Stoecklein NH, Heitzer E, Gironella M. Circulating biomarkers for early detection and clinical management of colorectal cancer. Mol Aspects Med 2019;69:107–22.

21. Alves Martins BA, de Bulhões GF, Cavalcanti IN, Martins MM, de Oliveira PG, Martins AMA. Biomarkers in colorectal cancer: The role of translational proteomics research. Front Oncol 2019;9:1284.

22. Cox J, Mann M. MaxQuant enables high peptide identification rates, individualized p.p.b.-range mass accuracies and proteome-wide protein quantification. Nat Biotechnol 2008;26:1367–72.

23. Bruderer R, Bernhardt OM, Gandhi T, Miladinović SM, Cheng LY, Messner S, Ehrenberger T, Zanotelli V, Butscheid Y, Escher C, Vitek O, Rinner O, Reiter L. Extending the limits of quantitative proteome profiling with data-independent acquisition and application to acetaminophen-treated three-dimensional liver microtissues. Mol Cell Proteomics 2015;14:1400–10.

24. Choi M, Chang CY, Clough T, Broudy D, Killeen T, MacLean B, Vitek O. MSstats: an R package for statistical analysis of quantitative mass spectrometry-based proteomic experiments. Bioinformatics. 2014;30:2524–6.

25. Yu G, Wang L-G, Han Y, He Q-Y. clusterProfiler: an R package for comparing biological themes among gene clusters. OMICS 2012;16:284–7.

26. Boyle EI, Weng S, Gollub J, Jin H, Botstein D, Cherry JM, Sherlock G. GO::TermFinder--open source software for accessing Gene Ontology information and finding significantly enriched Gene Ontology terms associated with a list of genes. Bioinformatics 2004;20:3710–5.

27. Subramanian A, Tamayo P, Mootha VK, Mukherjee S, Ebert BL, Gillette MA, Paulovich A, Pomeroy SL, Golub TR, Lander ES, Mesirov JP. Gene set enrichment analysis: a knowledge-based approach for interpreting genome-wide expression profiles. Proc Natl Acad Sci U S A 2005;102:15545–50.

28. Horton P, Park KJ, Obayashi T, Fujita N, Harada H, Adams-Collier CJ, Nakai K. WoLF PSORT: protein localization predictor. Nucleic Acids Res 2007;35:W585–W7.

29. Szklarczyk D, Gable AL, Lyon D, Junge A, Wyder S, Huerta-Cepas J, Simonovic M, Doncheva NT, Morris JH, Bork P, Jensen LJ, von Mering C. STRING v11: proteinprotein association networks with increased coverage, supporting functional discovery in genome-wide experimental datasets. Nucleic Acids Res 2019;47:D607–d13.

30. MacLean B, Tomazela DM, Shulman N, Chambers M, Finney GL, Frewen B, Kern R, Tabb DL, Liebler DC, MacCoss MJ. Skyline: an open source document editor for creating and analyzing targeted proteomics experiments. Bioinformatics 2010;26:966–8.

31. Metz CE. Basic principles of ROC analysis. Semin Nucl Med 1978;8:283–98.

32. Robin X, Turck N, Hainard A, Tiberti N, Lisacek F, Sanchez JC, Muller M. pROC: an open-source package for R and S+ to analyze and compare ROC curves. BMC Bioinformatics 2011;12:77.

33. Zhang B, Wang J, Wang X, Zhu J, Liu Q, Shi Z, Chambers MC, Zimmerman LJ, Shaddox KF, Kim S, Davies SR, Wang S, Wang P, Kinsinger CR, Rivers RC, Rodriguez H, Townsend RR, Ellis MJ, Carr SA, Tabb DL, Coffey RJ, Slebos RJ, Liebler DC; NCI CPTAC. Proteogenomic characterization of human colon and rectal cancer. Nature 2014;513:382–7.

34. Kleinschmidt EG, Miller NLG, Ozmadenci D, Tancioni I, Osterman CD, Barrie AM, Taylor KN, Ye A, Jiang S, Connolly DC, Stupack DG, Schlaepfer DD. Rgnef promotes ovarian tumor progression and confers protection from oxidative stress. Oncogene. 2019;38:6323–37.

35. Li MY, Peng WH, Wu CH, Chang YM, Lin YL, Chang GD, Wu HC, Chen GC. PTPN3 suppresses lung cancer cell invasiveness by counteracting Src-mediated DAAM1 activation and actin polymerization. Oncogene 2019;38:7002–16.

36. Zeng Y, Cao Y, Liu L, Zhao J, Zhang T, Xiao L, Jia M, Tian Q, Yu H, Chen S, Cai Y. SEPT9_i1 regulates human breast cancer cell motility through cytoskeletal and RhoA/FAK signaling pathway regulation. Cell Death Dis 2019;10:720.

37. Ailiken G, Kitamura K, Hoshino T, Satoh M, Tanaka N, Minamoto T, Rahmutulla B, Kobayashi S, Kano M, Tanaka T, Kaneda A, Nomura F, Matsubara H, Matsushita K. Post-transcriptional regulation of BRG1 by FIRΔexon2 in gastric cancer. Oncogenesis 2020;9:26.

38. Wu H, Zhang J, Dai R, Xu J, Feng H. Transferrin receptor-1 and VEGF are prognostic factors for osteosarcoma. J Orthop Surg Res 2019;14:296.

39. Shigeta S, Toyoshima M, Kitatani K, Ishibashi M, Usui T, Yaegashi N. Transferrin facilitates the formation of DNA double-strand breaks via transferrin receptor 1: the possible involvement of transferrin in carcinogenesis of high-grade serous ovarian cancer. Oncogene 2016;35:3577–86.

40. Shen Y, Li X, Dong D, Zhang B, Xue Y, Shang P. Transferrin receptor 1 in cancer: a new sight for cancer therapy. Am J Cancer Res 2018;8:916–31.

41. Neiveyans M, Melhem R, Arnoult C, Bourquard T, Jarlier M, Busson M, Laroche A, Cerutti M, Pugnière M, Ternant D, Gaborit N, Chardès T, Poupon A, Gouilleux-Gruart V, Pèlegrin A, Poul MA. A recycling anti-transferrin receptor-1 monoclonal antibody as an efficient therapy for erythroleukemia through target up-regulation and antibody-dependent cytotoxic effector functions. MAbs 2019;11:593–605.

42. Nakamura Y, Nakamichi N, Takarada T, Ogita K, Yoneda Y. Transferrin receptor-1 suppresses neurite outgrowth in neuroblastoma Neuro2A cells. Neurochem Int 2012;60:448–57.

43. Leoh LS, Kim YK, Candelaria PV, Martínez-Maza O, Daniels-Wells TR, Penichet ML. Efficacy and mechanism of antitumor activity of an antibody targeting transferrin receptor 1 in mouse models of human multiple myeloma. J Immunol 2018;200:3485–94.

44. Jamnongkan W, Thanan R, Techasen A, Namwat N, Loilome W, Intarawichian P, Titapun A, Yongvanit P. Upregulation of transferrin receptor-1 induces cholangiocarcinoma progression via induction of labile iron pool. Tumour Biol 2017;39:1010428317717655.

45. Daniels-Wells TR, Penichet ML. Transferrin receptor 1: a target for antibody-mediated cancer therapy. Immunotherapy 2016;8:991–4.

46. Adachi M, Kai K, Yamaji K, Ide T, Noshiro H, Kawaguchi A, Aishima S. Transferrin receptor 1 overexpression is associated with tumour de-differentiation and acts as a potential prognostic indicator of hepatocellular carcinoma. Histopathology. 2019;75:63–73.

47. Li Q, Mao L, Wang R, Zhu L, Xue L. Overexpression of S-adenosylhomocysteine hydrolase (SAHH) in esophageal squamous cell carcinoma (ESCC) cell lines: effects on apoptosis, migration and adhesion of cells. Mol Biol Rep 2014;41:2409–17.

48. Jones JJ, Wilcox BE, Benz RW, Babbar N, Boragine G, Burrell T, Christie EB, Croner LJ, Cun P, Dillon R, Kairs SN, Kao A, Preston R, Schreckengaust SR, Skor H, Smith WF, You J, Hillis WD, Agus DB, Blume JE. A plasma-based protein marker panel for colorectal cancer detection identified by multiplex targeted mass spectrometry. Clin Colorectal Cancer 2016;15:186–94.e13.

