## Supplementary figures and images for "Proteomics study of colorectal cancer and adenomatous polyps identifies TFR1, SAHH, and HV307 as potential biomarkers for screening"

### Fig.S1.tif

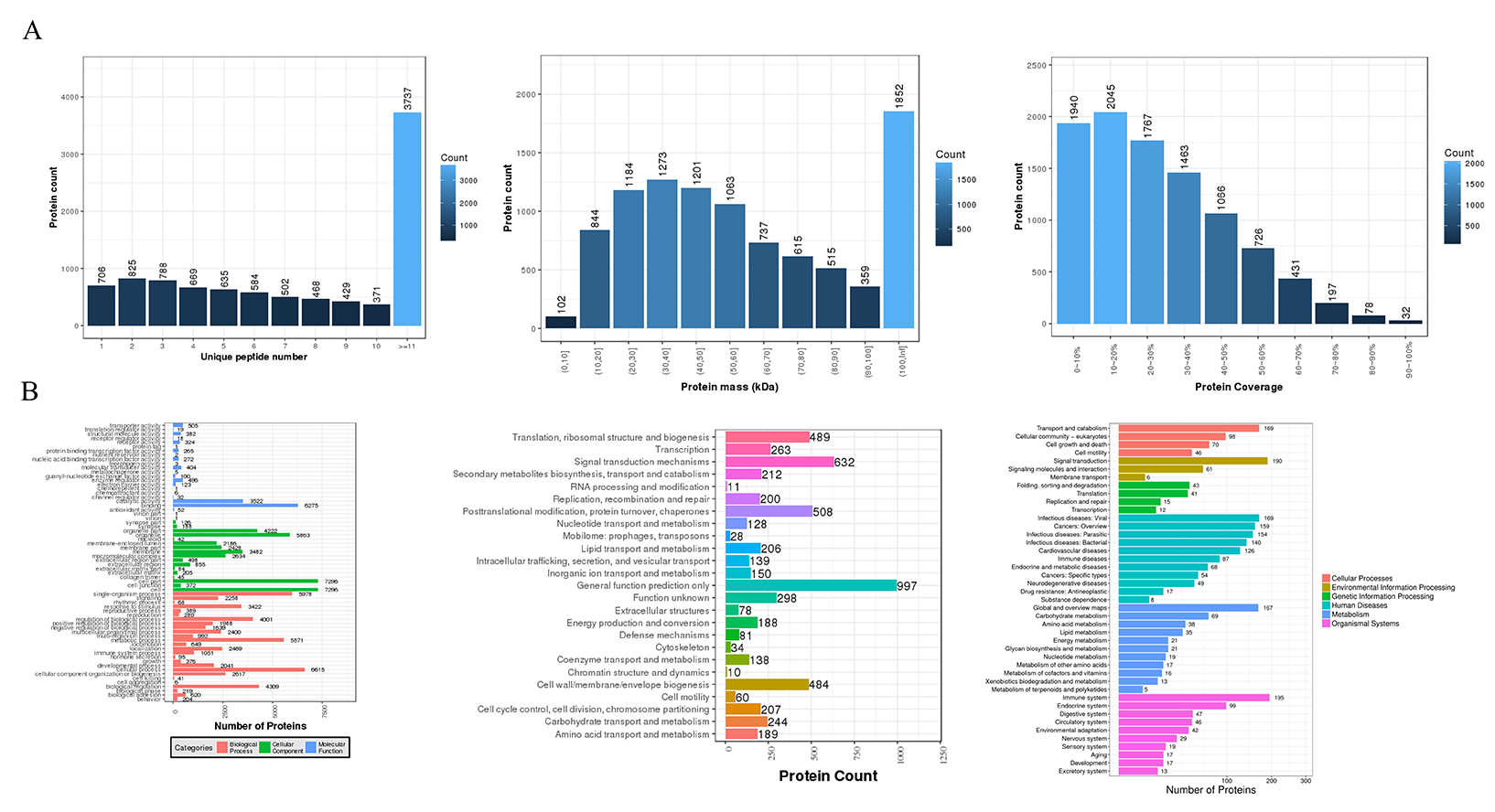

### Fig.S2.tif

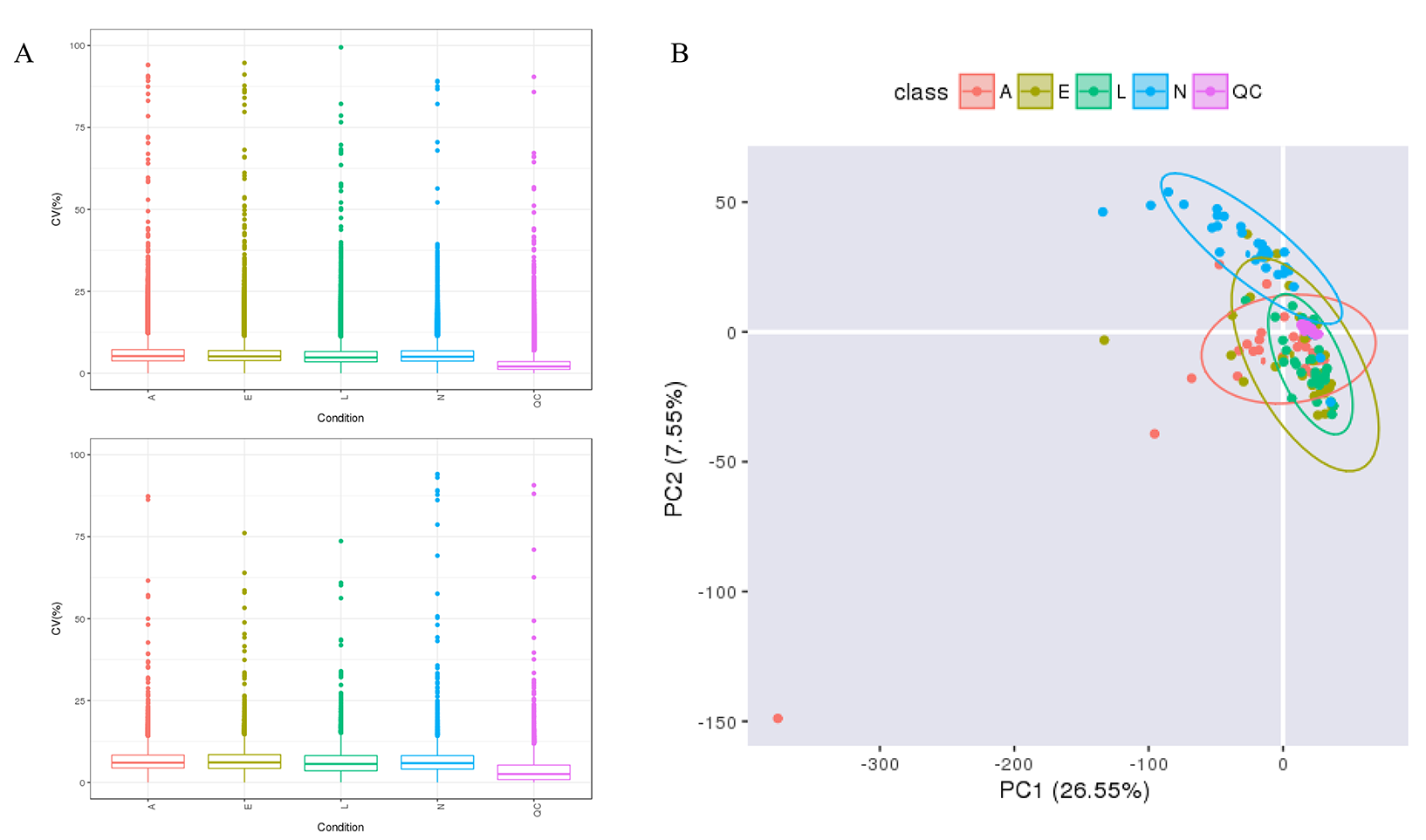

### Fig.S3.tif

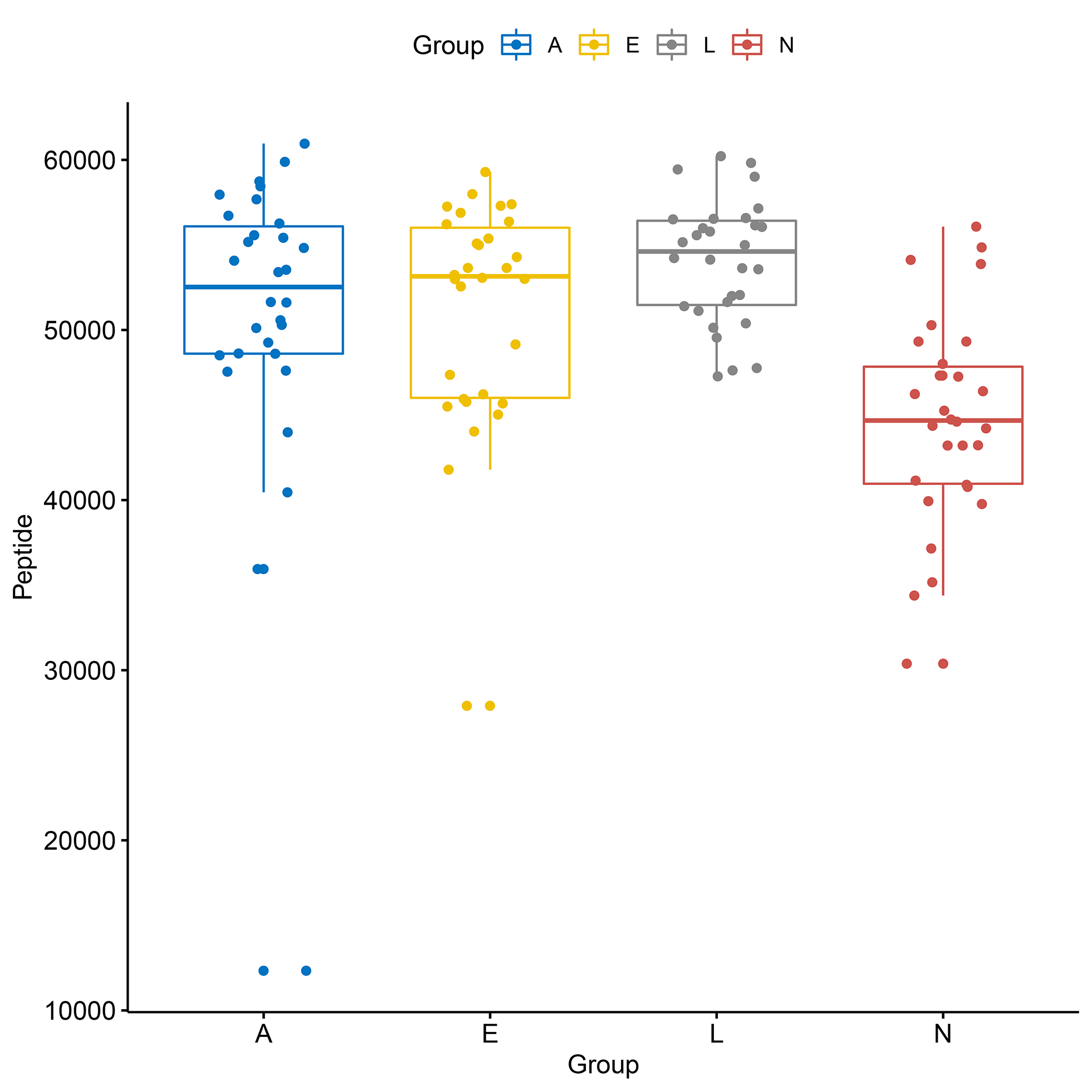

### Fig.S4.tif

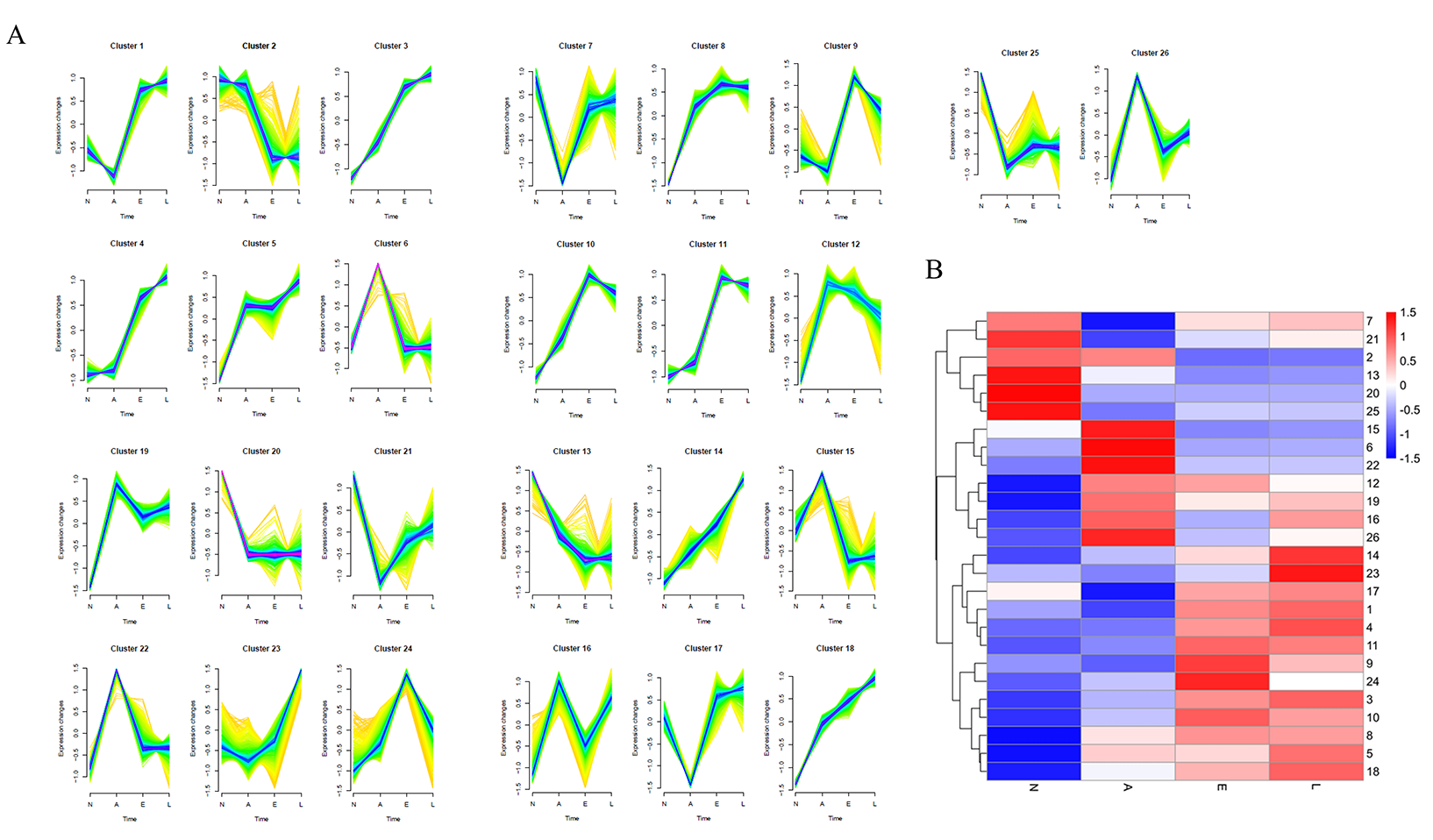

### Fig.S5.tif

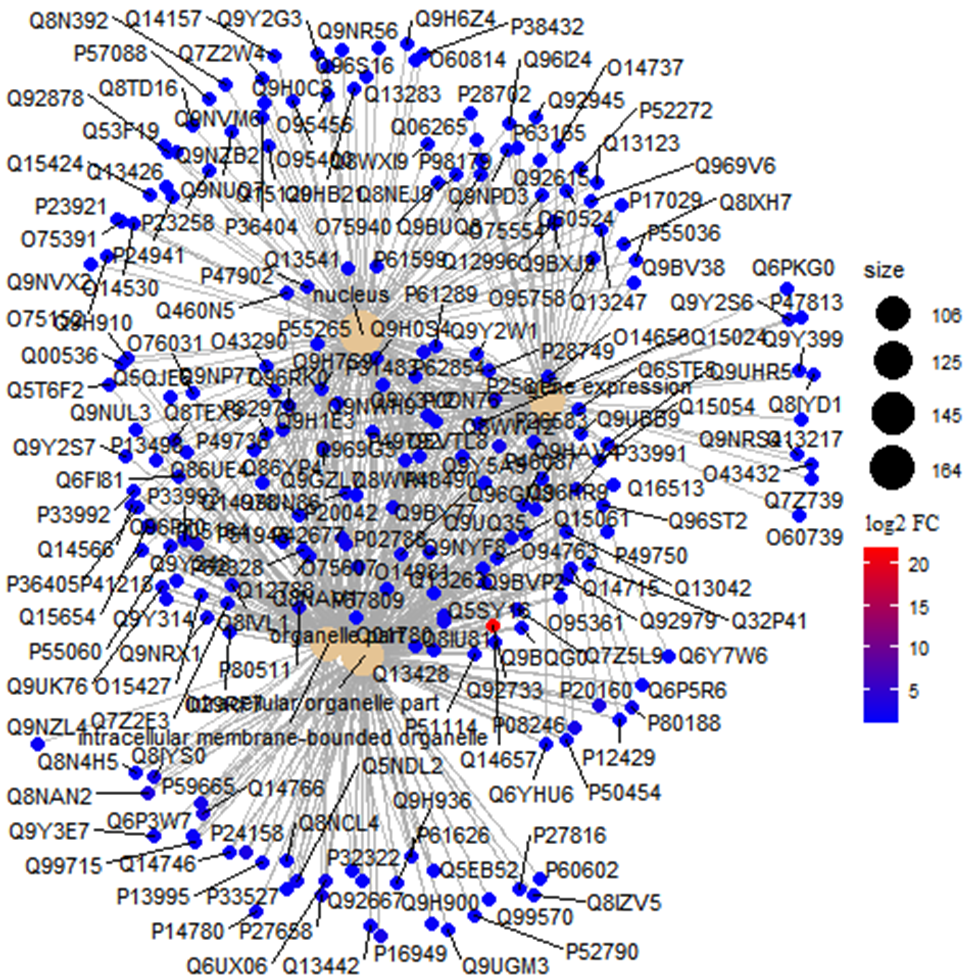

### Fig.S6.tif

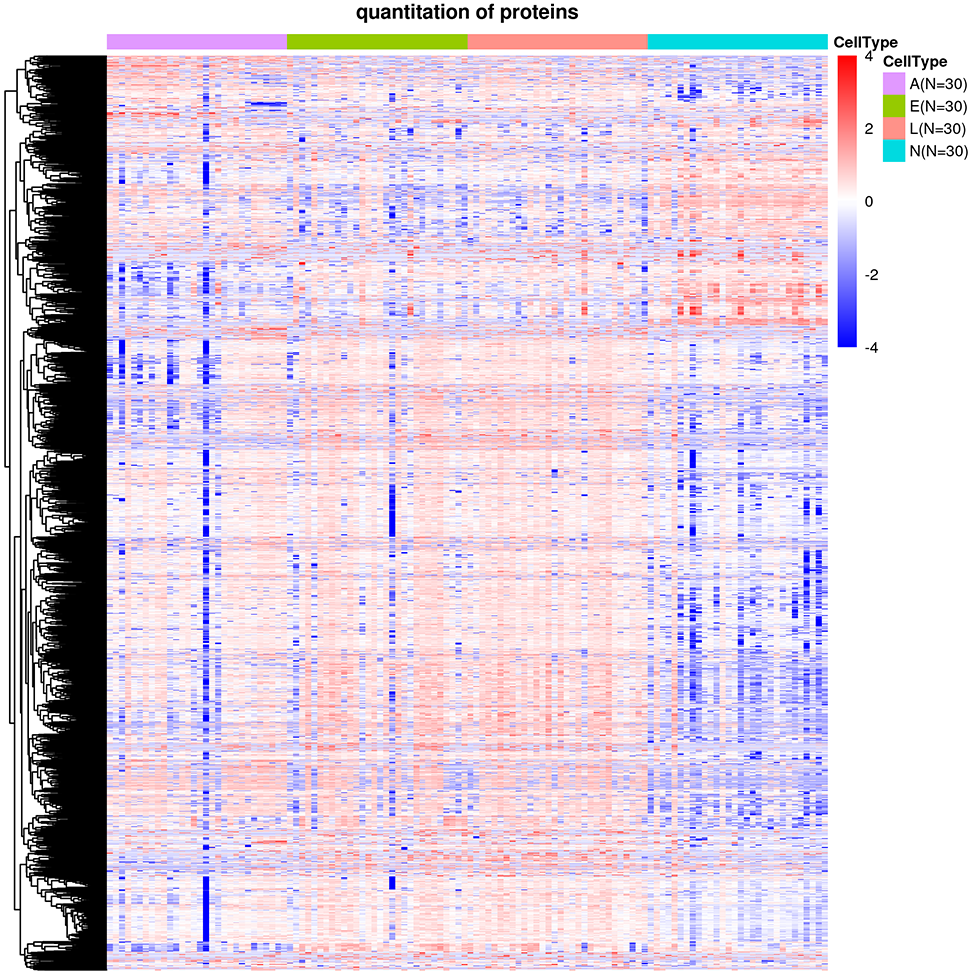

### Fig.S7.tif

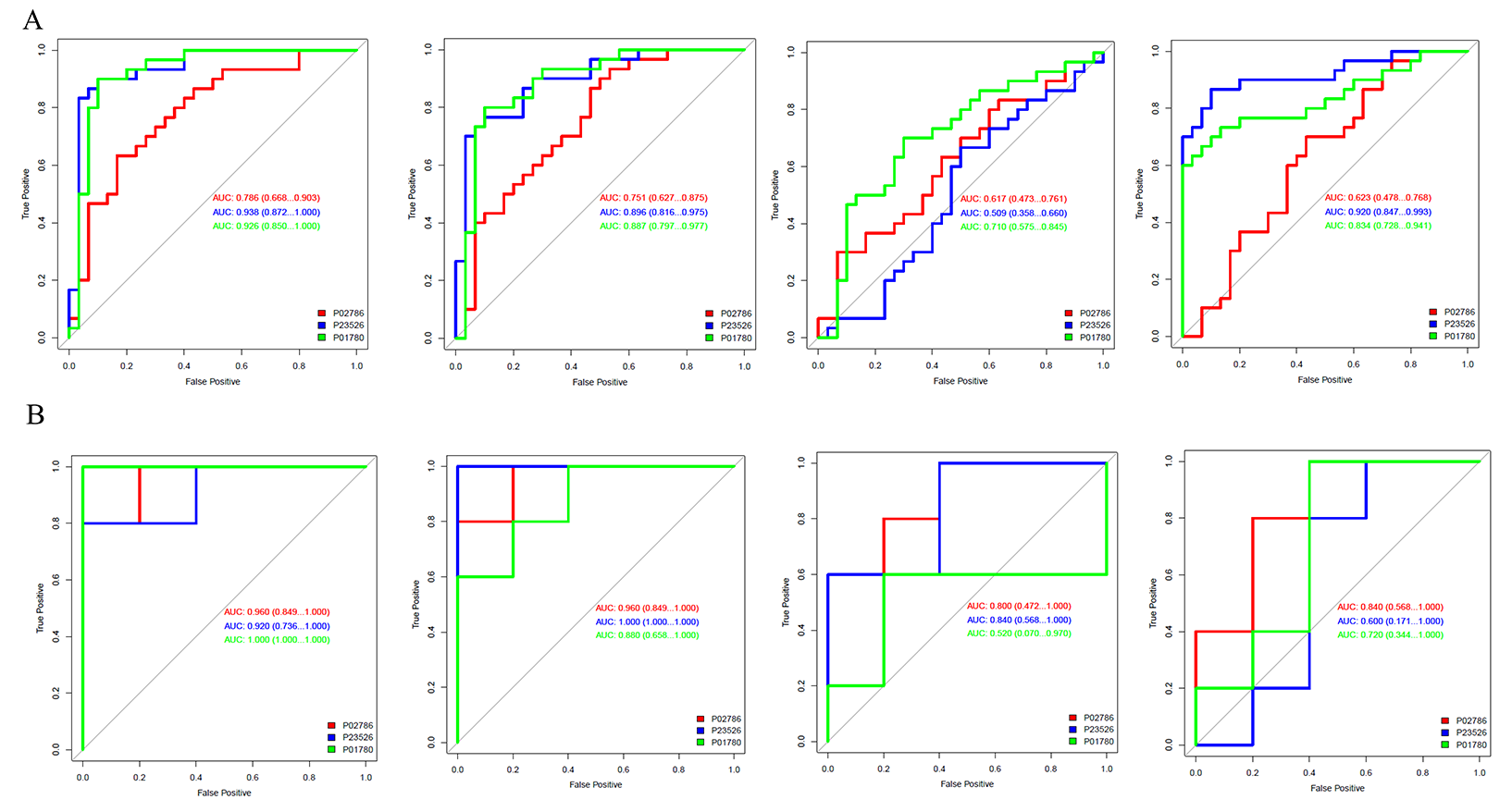
